# Interaction-Range Control of Synapsin Aggregation in a Coarse-Grained Model

**DOI:** 10.64898/2026.08.16.745062

**Authors:** Leandro B. Krott, Thiago Puccinelli, Walas de Oliveira, Maria E. N. Gomes, Enrique Lomba, Francesco Piazza, José Rafael Bordin

## Abstract

Synapsin-1 is a multidomain neuronal protein containing extensive intrinsically disordered regions and is a key component of synaptic-vesicle condensates. Direct residue-level simulation of the collective organization of thousands of synapsin molecules remains computationally demanding. Here, we develop a coarse-grained description that connects residue-level CALVADOS 3 simulations to a one-particle-per-protein model. A potential of mean force between two synapsin molecules is obtained by umbrella sampling and represented by an isotropic effective interaction containing a short-range attractive region and a weak outer repulsive contribution. We compare two treatments of this interaction that differ only in the retention of the outer tail. Langevin dynamics simulations of effective proteins show aggregation upon cooling and compression in both models, but with markedly different collective organization. The shorter-ranged model progressively coarsens toward a single dense domain, whereas retaining the outer repulsive contribution favors the persistence of multiple mesoscale aggregates. The two models also display distinct relationships between aggregate size and particle mobility at low temperature. These results show that weak features of an effective protein–protein interaction can have pronounced consequences for collective synapsin organization at mesoscopic scales.

**TOC Graphic:** 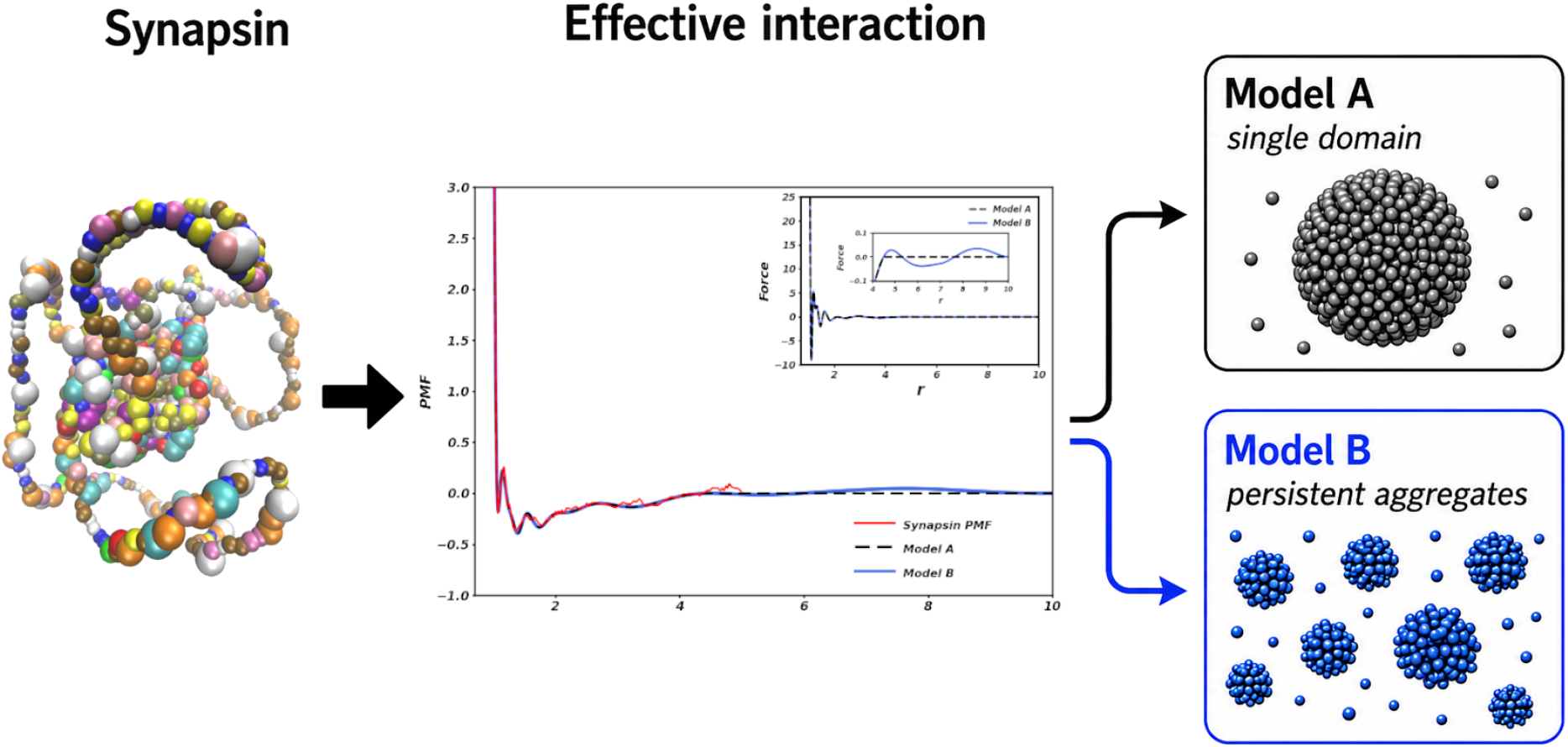

## Introduction

Biomolecular condensates provide a physical route for organizing cellular matter without delimiting membranes. Their formation and material properties emerge from a hierarchy of interactions that spans residue-scale contacts, polymer conformations, intermolecular association, interfacial organization, and mesoscale transport. This hierarchy is central to function: sequence-dependent interactions determine molecular partitioning and contact lifetimes, while the collective consequences are expressed through condensate growth, internal mobility, viscoelasticity, and exchange with the surrounding phase. Recent experimental and computational studies have emphasized that nanoscale interaction dynamics can propagate quantitatively into diffusion and bulk rheological response, underscoring the need for models that connect molecular resolution to collective behavior.^1–4^

Synaptic-vesicle clusters are a particularly rich example of this multiscale organization. Synapsins are abundant neuronal phosphoproteins associated with synaptic vesicles and have long been known to regulate the maintenance of the reserve pool required for sustained neurotransmitter release. The discovery that synapsin-1 can form a liquid phase that recruits lipid vesicles provided a condensate-based interpretation of vesicle clustering at presynaptic terminals.^5^ Subsequent work showed that a synapsin region capable of phase separation is required for vesicle clustering in a living synapse, and that coexpression of synapsin and synaptophysin is sufficient to generate liquid-like synaptic-vesicle-like clusters in nonneuronal cells.^6,7^ Synapsin condensates can also recruit other synaptic components, including *α*-synuclein, and can organize distinct vesicle populations into separate subphases, illustrating that the synapsin-rich environment is not merely an inert storage compartment.^8,9^

More recent experiments have strengthened the functional connection between condensation and synaptic-vesicle organization. Single-particle and super-resolution measurements showed that synapsin condensation controls vesicle sequestration, exchange, and mobility, linking condensate formation directly to vesicle dynamics. ^10^ Structural studies of reconstituted synapsin–vesicle condensates further revealed noncompact morphologies, close intervesicle contacts, and adhesion-induced vesicle deformation, while the aggregate size and geometry depend on protein concentration and the protein-to-lipid ratio.^11,12^ Measurements of the material response have added another level to this picture. Live-cell microrheology revealed a remarkably broad range of synapsin-condensate viscoelasticity that can be shifted by the partitioning of *α*-synuclein, whereas passive X-ray microrheology resolved distinct liquid-like and network-like regimes in synapsin-induced vesicle condensates.^13,14^ In addition, synapsin-1–vesicle condensates were recently shown to sequester actin and promote its polymerization, connecting the condensate to cytoskeletal organization and its regulation by phosphorylation.^15^ Collectively, these findings establish synapsin condensation as a dynamic organizing principle whose consequences extend from protein association to vesicle mobility, mesoscale morphology, rheology, and actin remodeling.

Despite this progress, the effective interactions responsible for the collective organization of large numbers of synapsin molecules remain incompletely characterized. Synapsin-1 is not a compact globular protein: it contains a folded central domain together with long intrinsically disordered terminal regions, generating a broad and anisotropic conformational ensemble. Recent mutational and biophysical analyses indicate that synapsin condensation is governed by sequence-encoded molecular grammars involving distributed interaction motifs rather than by a single binding site.^16^ Its intermolecular association therefore combines excluded volume, electrostatics, hydrophobic and sequence-specific contacts, conformational rearrangement, and solvent-mediated contributions. At the level of a protein-rich assembly, these microscopic contributions are integrated over many orientations and conformations, and weak features of the resulting effective interaction may influence whether dense domains coalesce, remain dispersed, or form long-lived finite aggregates. Isolating the consequences of the synapsin–synapsin interaction is thus a useful first step before the additional complexity of explicit vesicles, phosphorylation states, cytoskeletal components, and active presynaptic processes is introduced.

Molecular simulation can address this problem only through a compromise between resolution and accessible scale. Atomistic descriptions are valuable for resolving local interactions but are restricted to comparatively small systems and short times. One-bead-per-residue models extend the accessible range and have become important tools for studying the conformational ensembles and phase behavior of intrinsically disordered proteins.^17,18^ CALVADOS 3 advances this strategy by providing a unified representation of disordered regions and folded domains and by enabling simulations of multidomain proteins and their phase separation.^19^ This resolution is appropriate for characterizing the heterogeneous conformational ensemble of synapsin and for obtaining an interaction free energy between two proteins. However, a system of 5000 full-length synapsin-1 molecules would contain more than 3.5 million protein beads, before accounting for the long trajectories required to follow cluster growth, domain rearrangement, and collective diffusion. Direct residue-level exploration of the full density–temperature grid considered here would therefore be computationally prohibitive.

We consequently adopt a hierarchical coarse-graining strategy. In the first level, CALVADOS 3 retains residue-scale heterogeneity and conformational disorder. In the second, an entire synapsin molecule is represented by one effective interaction center, and the pair interaction is derived from the potential of mean force (PMF) between two residue-level proteins. This construction is bottom-up in the sense that the mesoscopic interaction inherits the free-energy landscape sampled by the higher-resolution model rather than being chosen solely to reproduce a prescribed macroscopic phase behavior. The PMF integrates over unresolved protein conformations, orientations, and implicit-solvent degrees of freedom at the reference condition, allowing the collective simulation to reach thousands of molecules and substantially longer length and time scales. Such a reduction is particularly useful for identifying generic relationships between the shape of an effective interaction and the resulting aggregation morphology. At the same time, it is necessarily an approximation: the PMF is state dependent, the one-center representation removes anisotropic and multivalent binding geometries, and pairwise additivity can become less accurate in a dense protein environment. The model should therefore be interpreted as a qualitative bridge across scales, not as a fully transferable molecular force field or a quantitative predictor of physiological coexistence conditions.^20^

This qualification is also central to the scientific question addressed here. The synapsin PMF contains a dominant short-range attraction together with a much weaker outer repulsive contribution. Because the latter is small in energy but extends over a larger volume, its collective importance cannot be inferred from the pair curve alone. We compare two representations that share the same short-distance interaction but differ in the treatment of this outer region: Model A truncates the interaction at an intermediate separation, whereas Model B smoothly retains the weak repulsive contribution over a longer range. This comparison provides a controlled test of how a seemingly minor choice made during coarse-graining and potential regularization propagates into aggregation, coalescence, morphology, pressure, and mobility at mesoscopic scales. Note that this long range repulsive tail turns the interaction of Model B into a Short Range Attractive-Long Range Repulsive (SALR) potential,^21–23^ a class of interactions that is known to stabilize cluster phases vs. plain two-phase demixing.^24^

Here, we first characterize the conformational heterogeneity of human synapsin-1 using CALVADOS 3 and obtain a two-protein PMF by umbrella sampling. We then construct the one-particle-per-protein models and perform Langevin dynamics simulations containing 5000 effective synapsin molecules over a broad range of reduced density and thermal energy. The resulting structures are analyzed through cluster statistics, representative morphologies, self-diffusion, and pressure. Our goal is not to assign a quantitatively predictive phase diagram to synapsin. Rather, we ask whether the interaction derived from a residue-level description is sufficient to generate collective aggregation and how retaining or removing its weak outer contribution changes the organization and coalescence of the protein-rich states. In this way, the study illustrates both the utility and the sensitivity of multiscale coarse-graining for biomolecular condensates.

## Models and Methods

### Residue-level model and single-chain characterization

The initial configuration of human synapsin-1 was obtained from the AlphaFold-predicted structure associated with UniProt entry P17600.^25^ The protein contains 705 residues. Following the structural assignment used here, residues 113–420 form the ordered central domain, while residues 1–112 and 421–705 constitute the disordered N- and C-terminal regions, respectively.^5^

Residue-level simulations were carried out at *T* = 293.15 K with the CALVADOS 3 coarse-grained model.^19^ The single-chain trajectory was used to characterize intramolecular contact probabilities, pair interaction energies, the radius of gyration *R_g_*, and the asphericity *Q*. These quantities provide a compact description of the spatial heterogeneity and conformational flexibility that are subsequently averaged into the mesoscopic pair interaction.

### Umbrella sampling and potential of mean force

The effective interaction between two synapsin molecules was determined by umbrella-sampling simulations performed with LAMMPS.^26^ The reaction coordinate was the distance *r* between the centers of mass of the ordered domains of the two proteins. A total of 88 overlapping harmonic windows were distributed along the reaction coordinate, with window centers ranging from 19.5 to 150.0 Å and separated by 1.5 Å. In each window, a harmonic biasing potential,

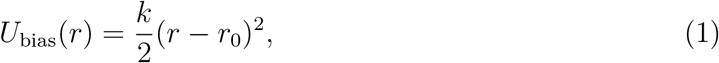

was applied, where *r*_0_ is the center of the umbrella window and *k* = 2 kcal mol*^−^*^1^ Å*^−^*^2^.Each simulation consisted of 5 × 10^6^ equilibration time steps followed by 5 × 10^6^ production time steps for data collection. The biased distributions were combined using the weighted histogram analysis method (WHAM),^27–32^ with 300 histogram bins. The iterative procedure was considered converged when the self-consistency tolerance reached 10*^−^*^8^.

The radial-volume contribution was subsequently removed from the free-energy profile obtained from WHAM by accounting for the 4*πr*^2^ Jacobian associated with the radial coordinate,

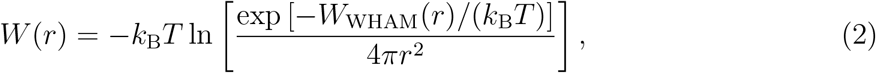

where *W*_WHAM_(*r*) denotes the profile obtained directly from WHAM and *T* = 293.15 K is the simulation temperature. The corrected PMF *W* (*r*) was then taken as the state-specific effective interaction between the two ordered-domain centers,

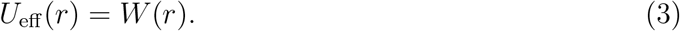

Because this PMF is obtained at one reference condition and along a single scalar coordinate, it averages over protein orientation and internal conformational degrees of freedom. It should consequently be interpreted as a reference effective interaction rather than as a temperature-independent microscopic potential.

### Analytical mesoscopic interaction

The PMF was represented by an analytical function combining a steep repulsive core, a broad attractive contribution, and Gaussian terms that reproduce intermediate-range wells and barriers,^33,34^

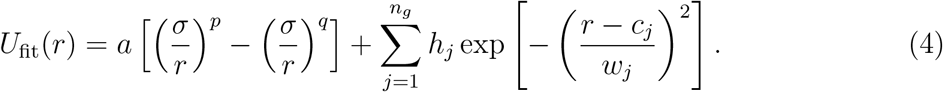

Here, *σ* = 30 Å defines the effective length scale and *ε* = 2.44 kJ mol*^−^*^1^ defines the energy scale. The fitted parameters are listed in Table 1.

**Table 1:** Parameters of the analytical effective interaction for human synapsin-1.

| Parameter | Value | Parameter | Value | Parameter | Value |
| --- | --- | --- | --- | --- | --- |
| $a$ | 25.2791 | $p$ | 10.4382 | $q$ | 3.38919 |
| $n_g$ | 6 | $\varepsilon$ | 2.44 kJ mol<sup>-1</sup> | $\sigma$ | 30 Å |
| $h_1$ | 3.49239 | $c_1$ | 1.34193 | $w_1$ | 0.276233 |
| $h_2$ | 3.68646 | $c_2$ | 1.20238 | $w_2$ | 0.155830 |
| $h_3$ | 2.06961 | $c_3$ | 1.55059 | $w_3$ | 0.484705 |
| $h_4$ | 3.20941 | $c_4$ | 1.10370 | $w_4$ | 0.104189 |
| $h_5$ | 0.14517 | $c_5$ | 3.77481 | $w_5$ | 1.143290 |
| $h_6$ | 1.34198 | $c_6$ | 1.87180 | $w_6$ | 1.016450 |

Two mesoscopic variants were used to probe the sensitivity to the outer interaction range. Model A employs the truncated and shifted potential

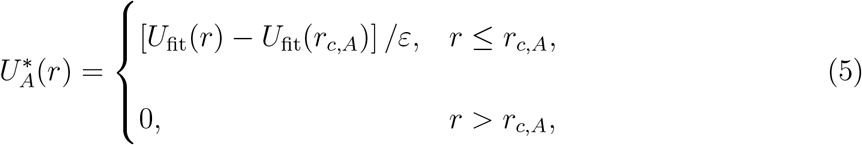

with *r_c,A_*= 4.5*σ*. Model B retains the analytical fitted interaction up to *r* = 7*σ* and extends it beyond this range using a quintic polynomial continuation up to *r_c,B_* = 10*σ*. The coefficients of the continuation are chosen such that the potential, its first derivative, and its second derivative match those of the fitted interaction at *r* = 7*σ*, while the potential and its first and second derivatives vanish at *r_c,B_* = 10*σ*. Thus, the short-distance part is shared by the two models, whereas Model A truncates the interaction at 4.5*σ*, while Model B retains the fitted outer contribution and smoothly brings it to zero between 7*σ* and 10*σ*. Since the umbrella-sampling data extend only up to *r* = 5*σ*, the fitted interaction between 5*σ* and 7*σ* and the subsequent quintic continuation beyond 7*σ* constitute controlled extrapolations of the effective interaction.

Beyond *r* = 7*σ*, where the analytical fit is used to represent the outer part of the interaction, Model B was smoothly extended to *r_c,B_*= 10*σ* using a quintic polynomial,

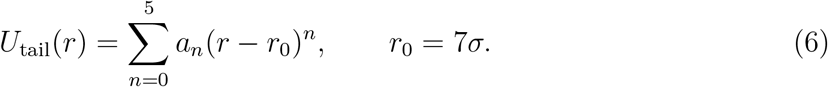

The six coefficients *a_n_* were determined by enforcing continuity of the potential and its first two derivatives at the connecting point,

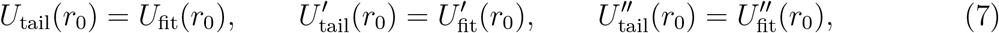

together with the conditions

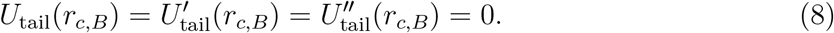

This construction provides a smooth *C*^2^ connection between the analytical fit and the zero-potential region.

Reduced quantities are defined as *r^∗^* = *r/σ*, *U ^∗^* = *U/ε*, *T ^∗^* = *k*_B_*T/ε*, and *ρ^∗^* = *ρσ*^3^. Asterisks are omitted below. Although *T* = 1 corresponds to the thermal energy of the PMF reference state, the temperature scan should be read as a variation of the ratio between thermal energy and the fixed effective interaction, not as a direct prediction over the corresponding range of physical temperatures.

### Mesoscopic simulations and observables

The collective simulations contained *N* = 5000 particles, each representing one synapsin molecule, in a cubic box with periodic boundary conditions. Molecular dynamics was performed in the canonical ensemble with LAMMPS using Langevin dynamics and damping parameter 1.0. The density was varied from *ρ* = 0.005 to 0.36169 and the reduced temperature from *T* = 3.00 to 1.00. At each density, the system was first equilibrated from a homogeneous fluid at *T* = 3.00 for 2 × 10^7^ steps, followed by 1 × 10^7^ production steps with time step Δ*t* = 0.001. The system was then cooled in increments of Δ*T* = 0.05, using the final configuration at the previous temperature as the initial state. Each subsequent temperature involved 10^6^ equilibration and 10^6^ production steps. Configurations, pressure, and energy were sampled every 2000 steps. Pressure was calculated from the virial expression.

Clusters were identified by a geometric connectivity criterion. Particles *i* and *j* were connected when

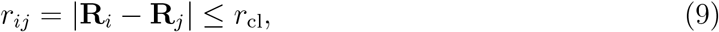

where *r*_cl_ = 1.4 was selected from the first-neighbor scale associated with the first minimum of the radial distribution function.^35–37^ Connected components of the resulting undirected graph were obtained using neighbor lists generated with the cKDTree implementation in SciPy. The number of connected components is denoted by *N*_clusters_. To suppress shortlived changes in graph membership, cluster assignments were filtered over ten consecutive stored configurations. This persistence criterion is used as a robust operational measure of aggregation and not as a thermodynamic order parameter.

Translational dynamics was characterized through the mean-squared displacement,

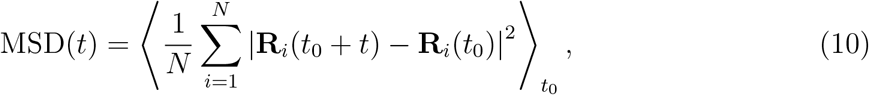

and the reduced self-diffusion coefficient was obtained from its long-time slope,

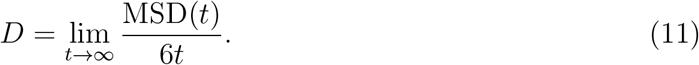

The absolute magnitude of *D* depends on the Langevin damping and is therefore used here to compare collective mobility across the simulated state points rather than to predict an experimental diffusion coefficient.

## Results and Discussion

Figure 1 summarizes the residue-level description of isolated synapsin-1. The contact-probability and pair-energy maps both reveal a compact central region embedded between much more heterogeneous terminal segments. The two maps are complementary: the contact probability reports how frequently residue pairs approach, whereas the energy map emphasizes whether those contacts are energetically favorable or unfavorable. Their common block-like central pattern is consistent with the ordered domain, while the diffuse off-diagonal features reflect contacts involving the disordered termini.

**Figure 1:**
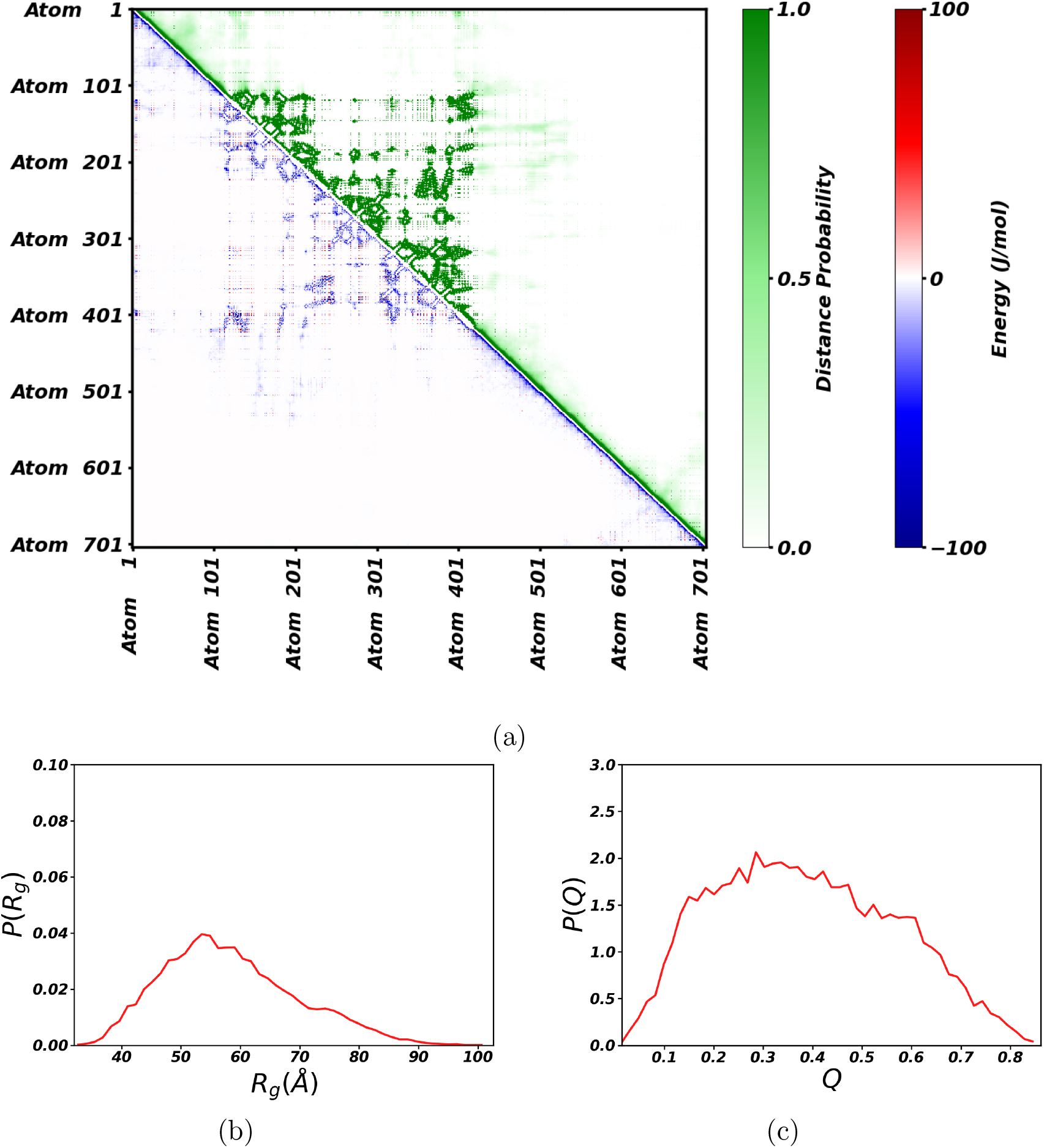
Residue-level characterization of human synapsin-1 from CALVADOS 3 simulations at 293.15 K. (a) Intramolecular distance-probability map (green, upper triangle) and pair interaction-energy map (blue/red, lower triangle). (b) Distribution of the radius of gyration *R_g_*. (c) Distribution of the asphericity *Q*.

The distribution of *R_g_* is broad, with its largest weight near 55 Å and a tail extending to substantially larger conformations. The asphericity distribution is likewise broad and centered at an intermediate value rather than close to zero. Synapsin therefore does not behave as a compact spherical object at the residue level. The one-particle mesoscopic description averages over an ensemble of extended and anisotropic conformations, which motivates treating the fitted interaction as an orientation-averaged free energy.

The PMF and its analytical representation are shown in Figure 2. The interaction contains a steep excluded-volume core, a short-range attractive minimum, and weaker oscillatory features at larger separation. Most importantly for the collective simulations, the fitted curve retains a broad, low-amplitude positive contribution outside the main attractive region. As mentioned, this combination is reminiscent of short-range-attraction/long-range-repulsion interactions, although the effective protein potential is softer and more structured than standard minimal SALR models.^35,36,38^

**Figure 2:**
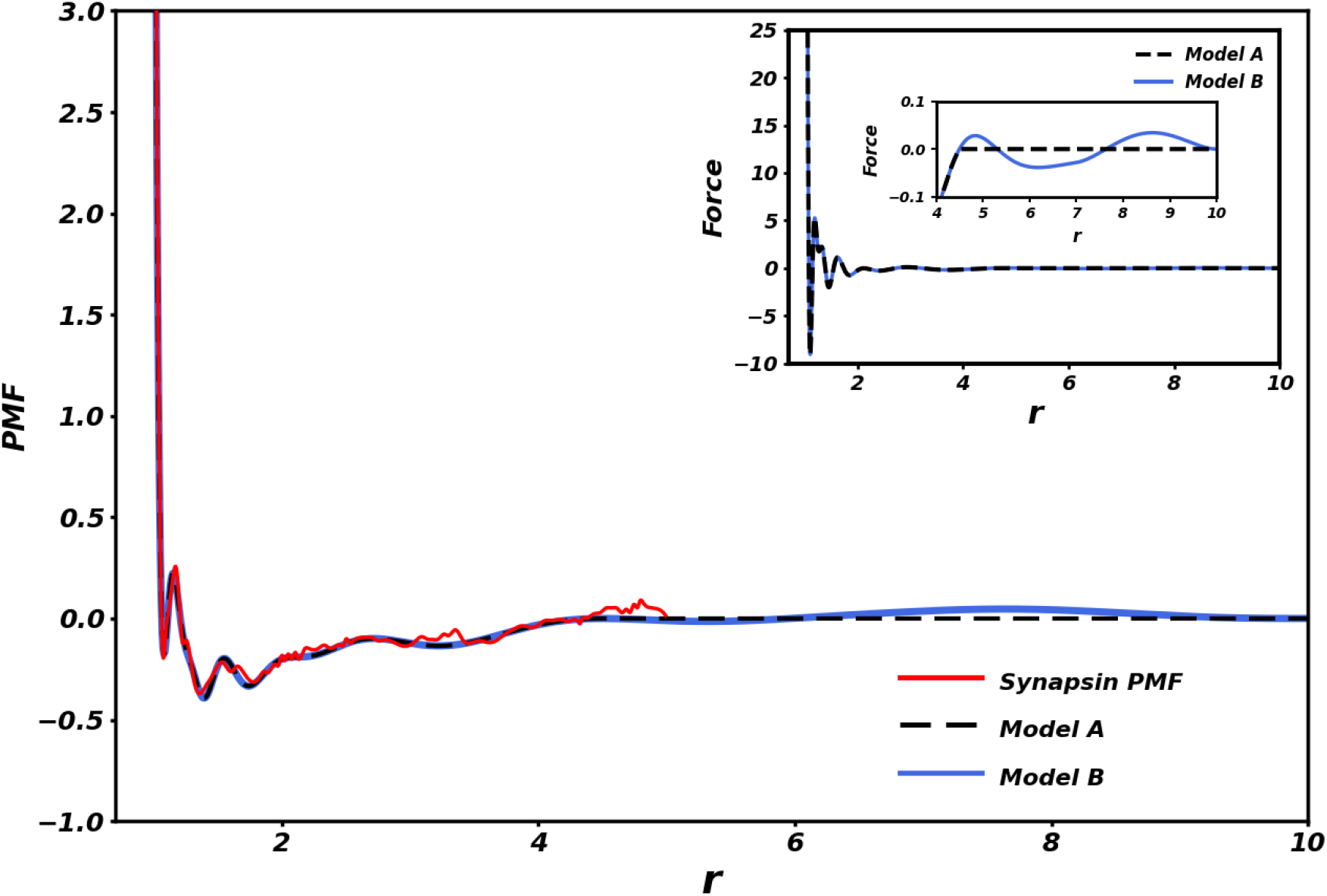
Potential of mean force for two human synapsin-1 molecules (red solid line), the truncated interaction potential (Model A, black dashed line), with *r_c,A_* = 4.5, and the analytically fitted interaction potential (Model B, blue solid line), extended to *r_c,B_* = 10. The inset shows the corresponding force profiles for Models A and B, obtained from the derivatives of their respective interaction potentials. The secondary inset provides a magnified view of the force profiles in the range 4 ≤ *r* ≤ 10, highlighting the differences introduced by the retained outer contribution in Model B. Distances and energies are reported in reduced units.

The two mesoscopic models isolate the effect of this outer contribution. Truncation at *r_c,A_*= 4.5 removes most of the broad repulsive tail, leaving an effectively shorter-ranged attraction. Extending the potential to *r_c,B_* = 10 retains that tail and adds a small free-energy cost for bringing distinct aggregates into closer proximity. Because the outer feature is weak relative to the short-distance interaction, its effect is not obvious from the pair potential alone; the collective simulations below show that it is nevertheless sufficient to reorganize aggregation pathways and final morphologies.

Figure 3 compares the temperature dependence of cluster connectivity and mobility. At low density and high temperature, *N*_clusters_*/N* is close to unity because most effective proteins are isolated or belong to very small connected components. Cooling promotes association and lowers this ratio. The change is particularly sharp for Model A at intermediate densities, consistent with rapid coarsening into one dominant connected aggregate. Model B displays the same overall tendency but generally retains a larger number of connected components, indicating that the outer repulsive contribution hinders complete merger.

**Figure 3:**
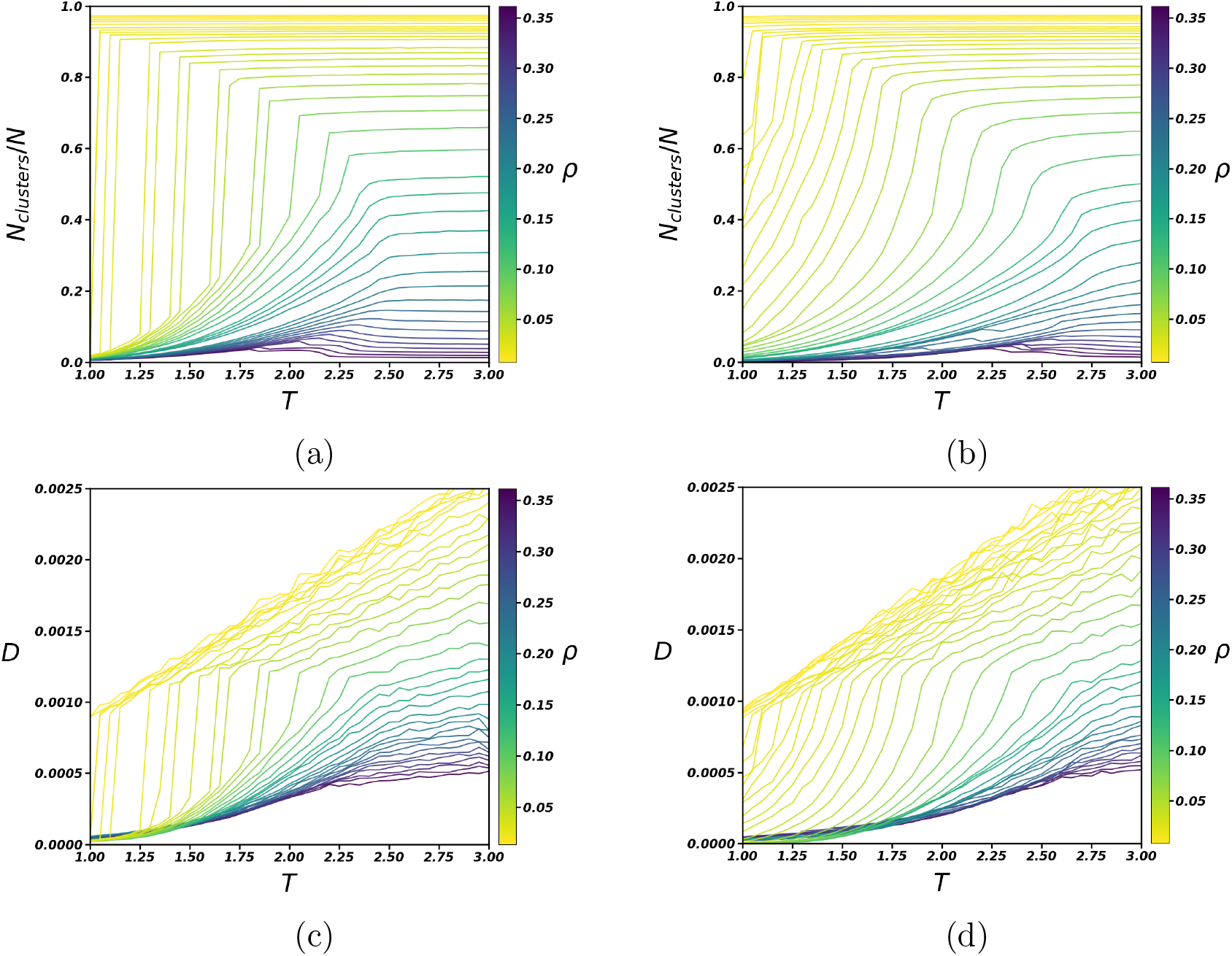
Temperature dependence of structural and dynamical observables. Normalized number of clusters, *N*_clusters_*/N*, for (a) Model A and (b) Model B. Reduced self-diffusion coefficient for (c) Model A and (d) Model B. Curves correspond to the densities indicated by the color bars.

The self-diffusion coefficient provides a complementary dynamical signature. At fixed density, *D* increases with temperature, whereas cooling and compression reduce translational mobility. The temperature interval in which the cluster ratio decreases is also the interval in which diffusion becomes strongly suppressed. This correspondence supports a collective crossover from a dispersed fluid to aggregated states. It does not by itself establish a thermodynamic binodal, because both observables are sensitive to connectivity, crowding, and the cooling history. We therefore use them to construct operational aggregation maps rather than equilibrium coexistence curves.

The same behavior is visible when the observables are plotted against density (Figure 4). At high temperature, compression produces a relatively smooth decrease of *N*_clusters_*/N* and of *D*. At lower temperature, the response becomes nonuniform and is accompanied by changes in the observed aggregate morphology. The symbols overlaid on the original curves identify morphological boundaries assigned from trajectory inspection and correlated changes in connectivity and mobility. In particular, the star shown for Model A marks the apparent terminal point of the sharp aggregation region in the present data set.

**Figure 4:**
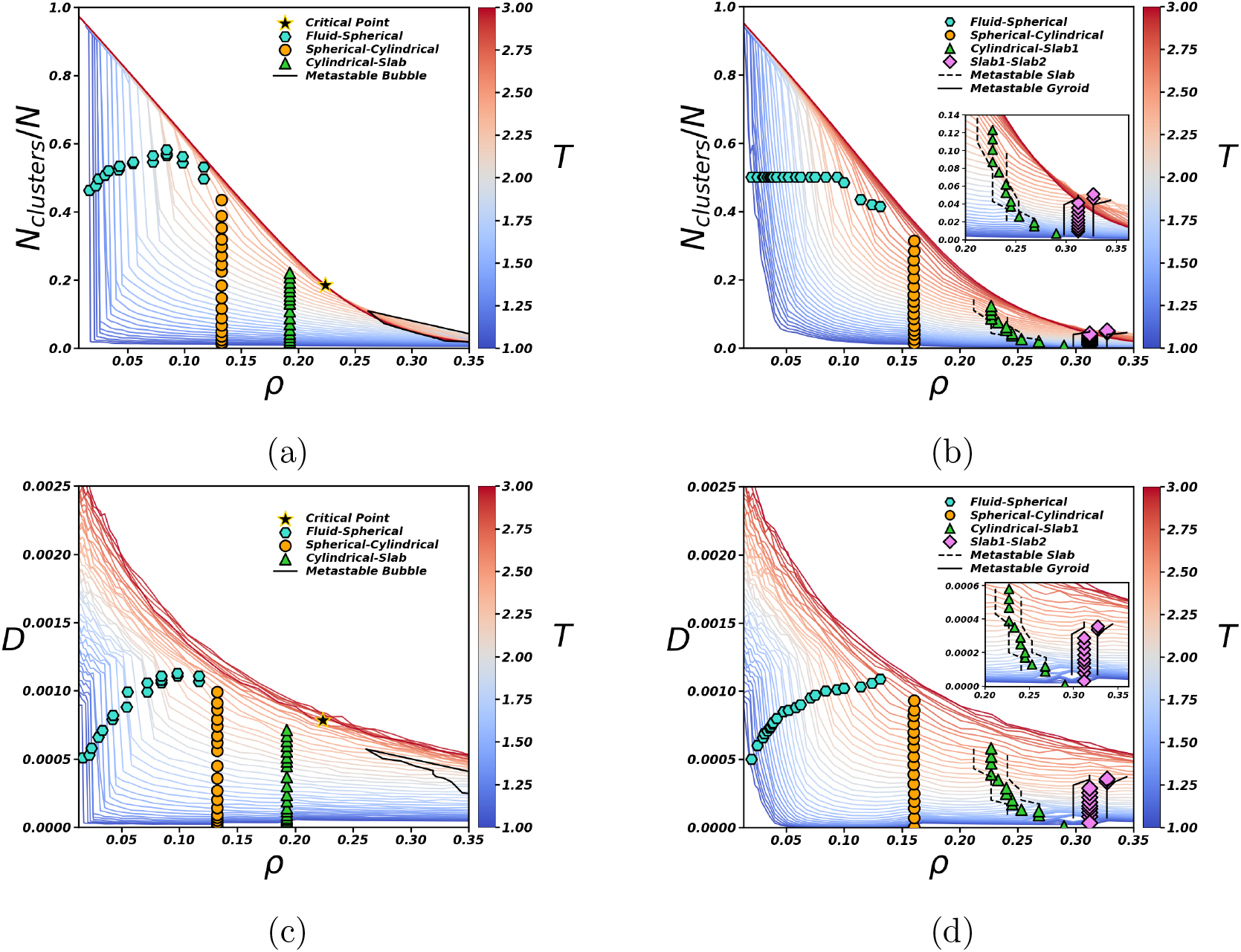
Density dependence of structural and dynamical observables. Normalized number of clusters for (a) Model A and (b) Model B, and reduced self-diffusion coefficient for (c) Model A and (d) Model B. Temperatures are indicated by the color bars. The overlaid symbols reproduce the trajectory-based morphological boundaries used in the state-map analysis. The star in Model A denotes an apparent endpoint of the aggregation region within the present finite-system protocol, not a finite-size-scaled critical point.

For Model A, the boundaries organize into a relatively compact region separating the dispersed fluid from single-domain morphologies. Model B exhibits a broader sequence of boundaries because multiple aggregates, elongated clusters, connected networks, and slablike arrangements occur over distinct density intervals. The comparison shows that the long-range tail changes more than the numerical location of aggregation: it changes how the dense material is partitioned throughout the simulation box.

To further resolve the dynamical consequences of aggregation, we compare the mobility of aggregate-associated and free synapsin particles in Figure 5. For aggregate-associated particles, the MSD is evaluated using the center of mass of the corresponding aggregate as reference, thereby removing its overall translational motion. To reduce the scatter associated with comparing states having similar aggregate sizes, the data at each temperature were grouped into 10 quantile bins in log_10_⟨*s*⟩, where ⟨*s*⟩ is the mean cluster size. Each point in Figure 5 represents the median ⟨*s*⟩ and the median ratio *D*_agg_*/D*_free_ within the corresponding bin. Only states with at least 30 samples in both the aggregate-associated and free populations were included. At *T* = 1.0, the two models show markedly different behavior. In Model A, *D*_agg_*/D*_free_ remains strongly below unity over most of the aggregated regime, consistent with the progressive coarsening toward a single dense domain. In Model B, the ratio instead increases systematically with aggregate size: mobility is strongly suppressed in small and intermediate aggregates, but approaches that of the free population as larger mesoscale aggregates develop. This behavior suggests that the weak longer-ranged repulsive contribution affects not only coalescence and morphology, but also the coupling between aggregate size and internal synapsin mobility.

**Figure 5:**
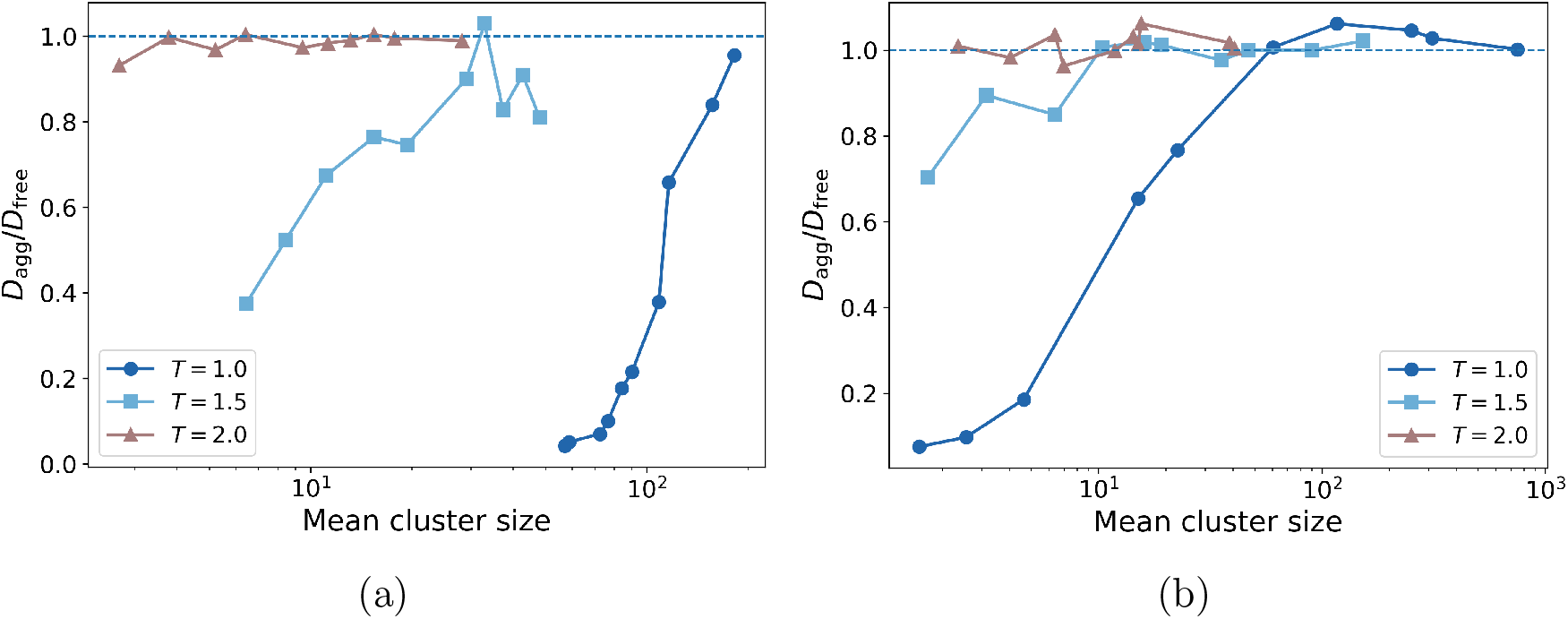
Relative mobility of aggregate-associated and free synapsin particles as a function of mean cluster size for (a) Model A and (b) Model B. *D*_agg_ is obtained from the MSD of aggregate-associated particles using the center of mass of the corresponding aggregate as reference, while *D*_free_ is obtained from the free-particle population. Data are shown for reduced temperatures *T* = 1.0, 1.5, and 2.0. The horizontal dashed line marks *D*_agg_ = *D*_free_.

The dynamical contrast becomes progressively weaker upon increasing temperature. At *T* = 1.5, reduced mobility within aggregates is still visible, particularly for smaller aggregates, whereas at *T* = 2.0 the two diffusivities are generally comparable in both models. This trend provides a qualitative connection with the single-molecule measurements of Hoffmann *et al.*, which showed that synapsin remains mobile within condensates while its motion is suppressed relative to less confined environments, with an additional reduction in mobility in the presence of synaptic vesicles. ^10^ It is also consistent with the broader picture of synaptic-vesicle clusters as dynamic liquid-like assemblies in which confinement coexists with molecular rearrangement.^39^ In this respect, Model B is particularly interesting because the longer-ranged repulsion prevents unrestricted coalescence and stabilizes multiple mesoscale aggregates, while still producing a pronounced dynamical confinement at low temperature for small and intermediate aggregate sizes. The comparison therefore indicates that weak longer-ranged interactions can simultaneously regulate aggregate persistence, size, and internal mobility.

The virial pressure provides an independent mechanical view of the same collective rearrangements (Figure 6). At high temperature, pressure increases smoothly with density in both models. At lower temperature, Model A develops branches with reduced slope and abrupt changes near the density intervals in which the trajectory changes from dispersed particles to droplet, cylinder, slab, and inverted geometries. Such nonmonotonic finite-system branches are expected to be sensitive to metastability, periodic geometry, and the sequential cooling protocol. We therefore do not apply a Maxwell construction or interpret the marked points as rigorously determined coexistence densities.

**Figure 6:**
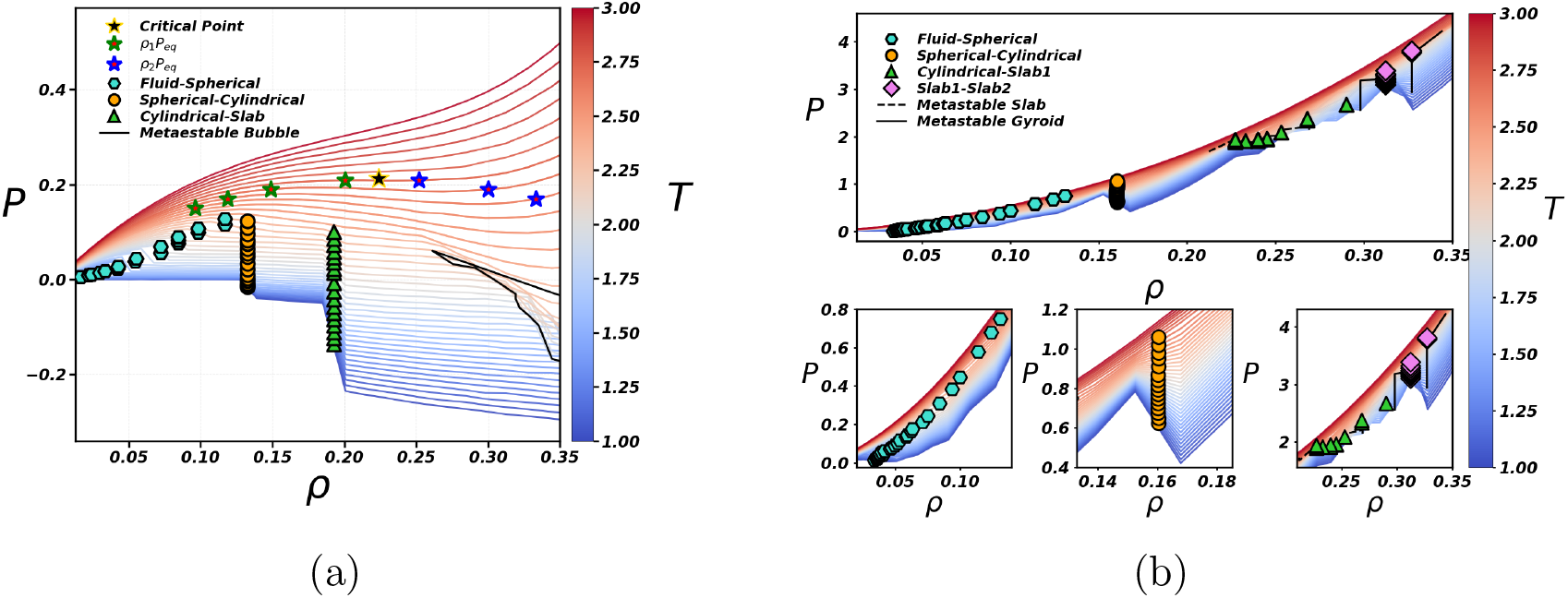
Virial pressure as a function of density for (a) Model A and (b) Model B. Tem-peratures are indicated by the color bars. Symbols reproduce the operational boundaries assigned from the trajectories and from correlated changes in cluster connectivity, diffusion, and pressure. The star in panel (a) is the apparent endpoint used in the original state-map analysis and is not interpreted here as a quantitatively determined critical point.

Model B retains a more strongly increasing pressure at high density, consistent with the additional outer repulsive contribution. Its lower-temperature curves also display slope changes correlated with the formation and reconnection of multiple aggregates. The agreement between pressure anomalies and the independently assigned structural boundaries supports the internal consistency of the state maps, while the different topology of the two maps demonstrates the collective impact of the interaction tail.

Representative Model A configurations at *T* = 1.85 are shown in Figure 7. At *ρ* = 0.0587, the system remains predominantly dispersed. A compact dense droplet is present at *ρ* = 0.0675, followed by an elongated cylindrical domain at *ρ* = 0.1526 and a slab-like structure at *ρ* = 0.201. At the highest density, *ρ* = 0.362, the dense region occupies most of the box and leaves a dilute cavity, producing an inverted or bubble-like morphology.

**Figure 7:**
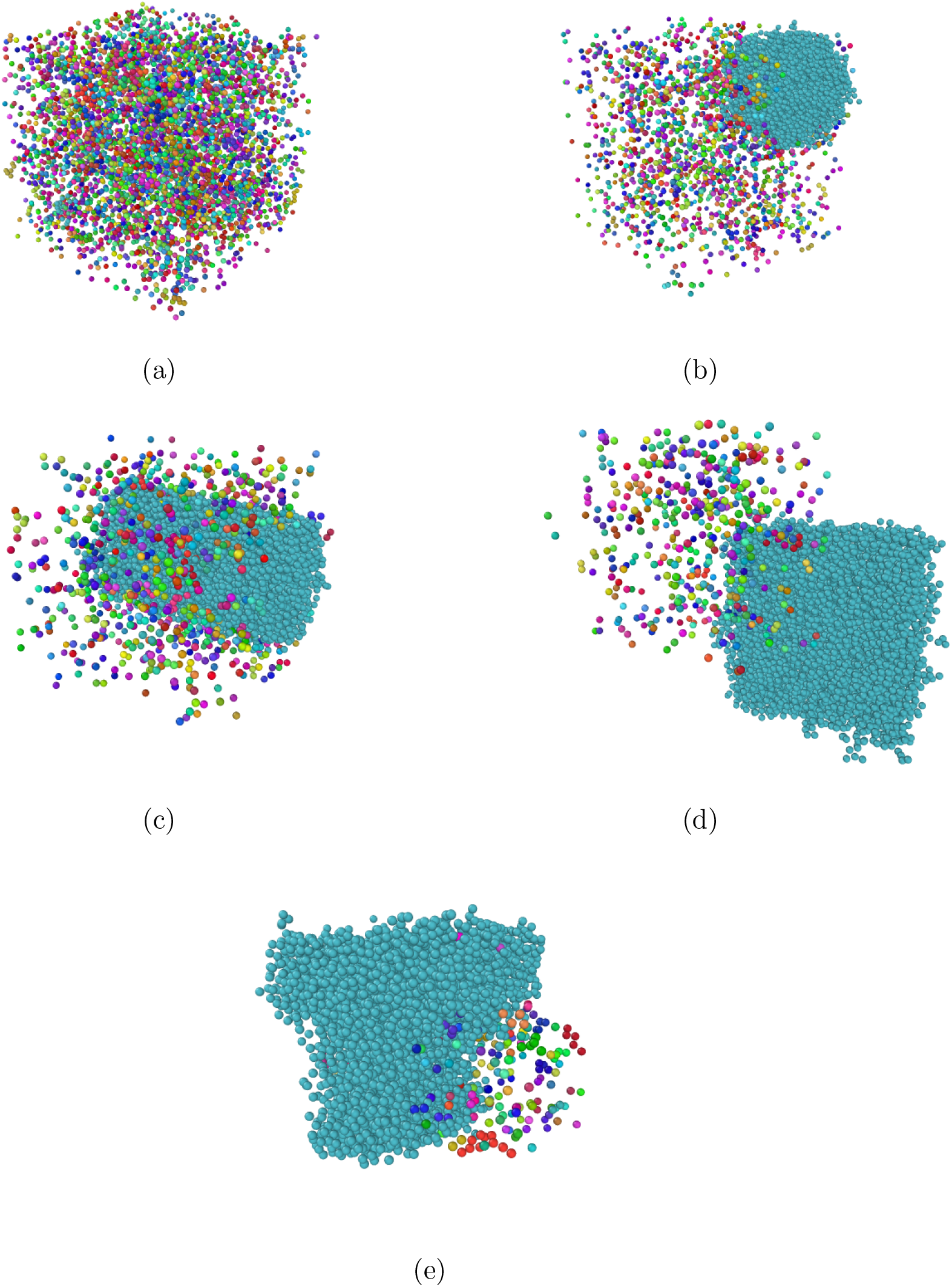
Representative unwrapped configurations of Model A at *T* = 1.85: (a) dispersed fluid at *ρ* = 0.0587; (b) one approximately spherical dense domain at *ρ* = 0.0675; (c) cylindrical domain at *ρ* = 0.1526; (d) slab-like domain at *ρ* = 0.201; and (e) inverted, bubble-like morphology at *ρ* = 0.362. Colors identify connected components in the cluster analysis; the dominant aggregate is shown in cyan. The shape sequence reflects finite-size and periodic-boundary effects within the aggregated region.

This droplet–cylinder–slab–inverted sequence should not be assigned to distinct bulk phases. In a periodic canonical box containing a fixed amount of dense and dilute material, the shape of the domain changes as the dense-phase volume fraction increases so as to reduce interfacial area. The snapshots nevertheless establish an important result: after crossing the aggregation region, Model A predominantly coarsens toward a single macroscopic connected domain. The remaining dilute particles and changes in domain shape account for part of the structure seen in the cluster, diffusion, and pressure curves.

Model B displays a qualitatively different organization at the same temperature (Figure 8). The low-density state at *ρ* = 0.0285 is dispersed. At *ρ* = 0.139, several compact droplets coexist instead of rapidly merging into one domain. Increasing the density produces multiple elongated aggregates at *ρ* = 0.228, followed by connected and partially percolating structures at *ρ* = 0.254. At still larger densities, slab-like and perforated morphologies appear, and more than one dense object may remain visible.

**Figure 8:**
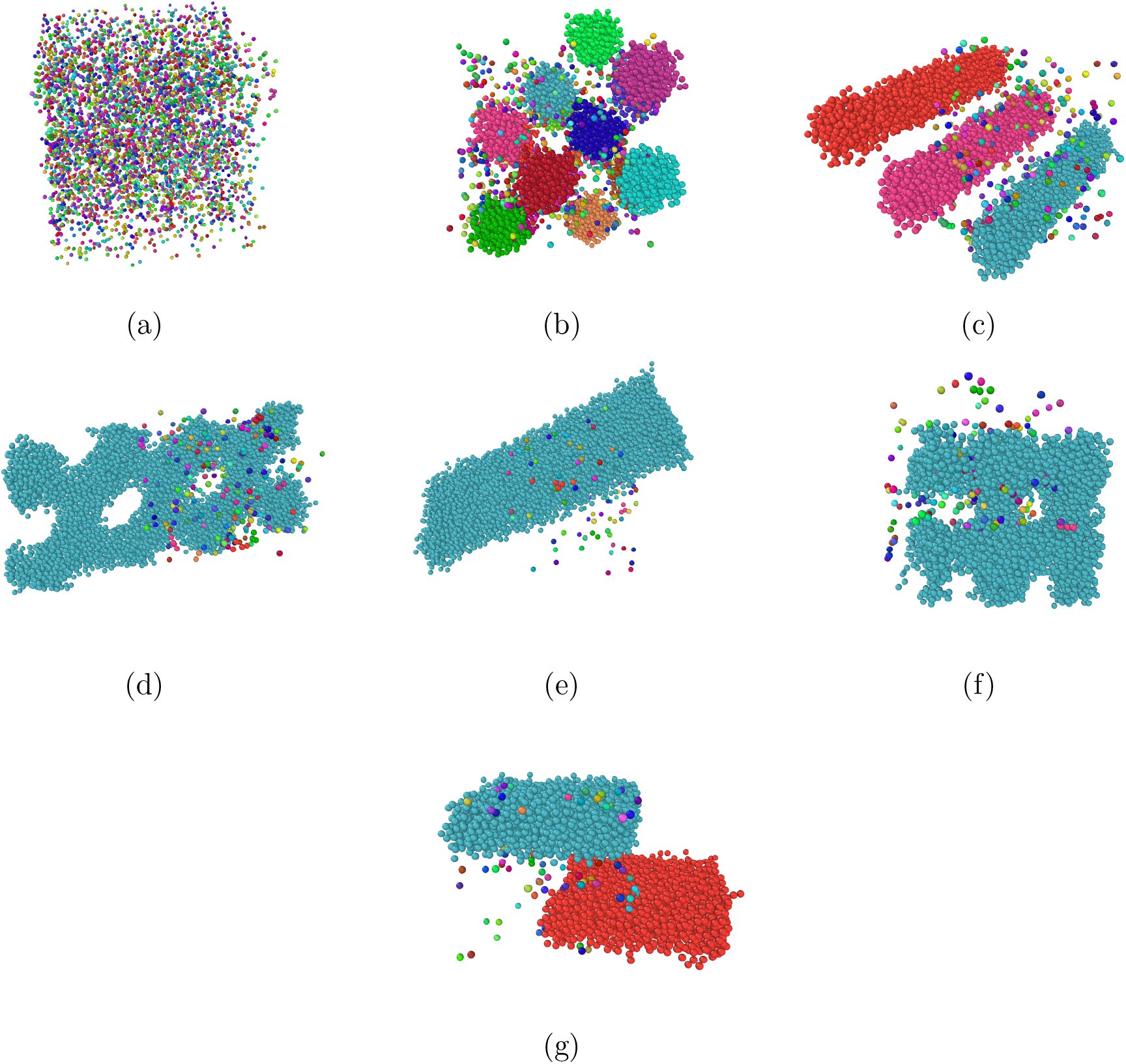
Representative unwrapped configurations of Model B at *T* = 1.85: (a) dispersed fluid at *ρ* = 0.0285; (b) multiple compact aggregates at *ρ* = 0.139; (c) multiple elongated aggregates at *ρ* = 0.228; (d) connected network-like morphology at *ρ* = 0.254; (e) slab-like structure at *ρ* = 0.298; (f) perforated connected morphology at *ρ* = 0.312; and (g) multiple slab-like dense domains at *ρ* = 0.362. Colors identify connected components. These structures are described as persistent morphologies over the simulated time window rather than as independently established equilibrium phases.

Within the simulated observation window, Model B sustains multiple finite-size aggregates over a broad range of conditions, contrasting to the progressive coalescence observed for Model A. This behavior is consistent with the competition characteristic of SALR-like interactions: the short-range attraction promotes local association, whereas the weak longer-ranged repulsion penalizes unrestricted domain growth and the merger of already formed dense regions.^36,38^**^?^**

## Conclusions

We developed a two-level coarse-grained description of human synapsin-1 in which residue-level CALVADOS 3 simulations and umbrella sampling provide a reference effective interaction for simulations of thousands of proteins. Cooling and compression produce aggregation together with correlated changes in cluster connectivity, translational mobility, and virial pressure.

The collective outcome is highly sensitive to the weak outer part of the effective interaction. The shorter-ranged Model A progressively coarsens toward a single dense domain, whereas Model B, which retains the weak longer-ranged repulsive contribution, sustains multiple mesoscale aggregates and inhibits unrestricted coalescence over the simulated time window. The dynamical analysis further shows that aggregate formation does not simply immobilize all associated proteins. At low temperature, aggregate-associated particles are substantially less mobile than the free population, while in Model B this dynamical contrast depends strongly on aggregate size and progressively weakens upon increasing temperature. This combination of persistent mesoscale organization and reduced but finite internal mobility provides a qualitative connection with experimental observations of synapsin-rich condensates as dynamic assemblies that constrain, rather than completely arrest, molecular motion. More generally, the results demonstrate that a bottom-up mesoscopic model can transfer molecular information to collective scales while revealing an important sensitivity: weak, long-ranged features of an effective protein–protein interaction can determine not only whether dense material coalesces, but also how it is partitioned and how molecules move within the resulting assemblies. Such features should therefore be considered explicitly when highly coarse-grained models are used to interpret biomolecular condensation.

## Acknowledgements

EL acknowledges the financial support from Grant No. PID2023-151751NB-I00 funded by MICIU/AEI/10.13039/501100011033 and by “ERDF A way of making Europe.” Galicia’s Supercomputing Center (CESGA) is also acknowledged for the generous allocation of computer resources. JRB and LBK thanks financial support from Brazilian National Council for Scientific and Technological Development (CNPq), grant n*^o^* 441728/2023-5. JRB also thanks CNPq, grant n*^o^* 309600/2026-0, Fundação de Amparo à Pesquisa do Estado do Rio Grande do Sul (FAPERGS), grant n*^o^* 25/2551-0002609-1, and Coordination for the Improvement of Higher Education Personnel (CAPES Finance Code 001) and the Alexander von Humboldt Foundation for financial support through a research fellowship at the University of Konstanz. TP thanks CAPES - Finance Code 001. MENG thanks Fundação de Amparo à Pesquisa do Estado de São Paulo (FAPESP), grant n*^o^* 2024/18313-0.

## Notes

### Competing Interest Statement

The authors have declared no competing interest.

